# LRIT3 is required for nyctalopin expression and normal ON and OFF pathway signaling in the retina

**DOI:** 10.1101/431338

**Authors:** Nazarul Hasan, Gobinda Pangeni, Thomas A. Ray, Kathryn M. Fransen, Jennifer Noel, Bart G. Borghuis, Maureen A. McCall, Ronald G. Gregg

**Author notes:** Corresponding Author: Ronald G. Gregg, Department of Biochemistry and Molecular Genetics, University of Louisville, 319 Abraham Flexner Way, Louisville, KY 40292.

## Abstract

At its first synapse, the retina establishes two parallel channels that encode light increments (ON) or decrements (OFF). At the same synapse, changes in photoreceptor glutamate release are sensed by ON bipolar cells (BCs) via the metabotropic glutamate receptor 6 (mGluR6), and OFF BCs via ionotropic BCs, which differ in their synaptic configuration with the photoreceptor terminal. ON BCs form invaginating synapses that bring them in close proximity to presynaptic ribbons and the presumed sole source of glutamate release. OFF bipolar cells form flat contacts distal to the ribbon synapse. We investigated the role of LRIT3 in normal assembly and function of the mGlur6 signaling cascade present in ON BCs. We demonstrate that LRIT3 is required for nyctalopin expression and thus TRPM1 expression and function. Using glutamate imaging, whole-cell electrophysiology, and multi-electrode array extracellular recordings we demonstrate that the loss of LRIT3 impacts both the ON and OFF pathways at the level of the BCs. The effect on ON pathway signaling, a lack of ON BC response, is shared by mutants lacking mGluR6, TRPM1 GPR179 or nyctalopin. The effects on the OFF pathway are unique to LRIT3, and include a decrease in response amplitude of both OFF BC and GCs. Based on these results, we propose a working model where LRIT3 is required for either efficient glutamate release or reuptake from the first retinal synapse.

**SIGNIFICANCE STATEMENT:** At the first visual synapse, photoreceptor cells signal to two distinct bipolar cell (BC) populations, one characterized by a depolarizing response to light onset (ON or DBCs), the other by a hyperpolarizing response (OFF or HBCs). The DBC light response depends on a G protein-coupled receptor and associated protein complex, known as the signalplex. Mutations in signalplex proteins lead to DBC pathway-specific loss of visual function. Here we show how loss of LRIT3, a previously identified signalplex protein, prevents functional assembly of the DBC signalplex and alters visual function in both ON and OFF signaling pathways. Thus, our results indicate that the function of LRIT3 at this first synapse extends beyond assembly of the DBC signalplex.

## INTRODUCTION

Light signaling is initiated in the retina when photoreceptors detect a luminance increase, hyperpolarize, and decrease tonic glutamate release. Several types of bipolar cells (BCs) detect this signal and establish several parallel information channels. First, the dichotomy of rod and cone photoreceptor light sensitivity creates channels that encode visual signals under dim and bright illumination, and signal through rod and cone BCs, respectively. Second, a difference in the glutamate receptors expressed by rod and cone ON bipolar cells (mGluR6; (Kaneko and Saito, 1983; Saito and Kaneko, 1983; Slaughter and Miller, 1983; Borghuis et al., 2014)) vs. OFF BCs (AMPA/kainate; (Kaneko and Saito, 1983; Saito and Kaneko, 1983; Slaughter and Miller, 1983; DeVries and Schwartz, 1999; Borghuis et al., 2014; Ichinose and Hellmer, 2016)) causes a light increment to depolarize ON BCs (DBCs) while hyperpolarizing OFF BCs (HBCs).

There are distinct differences in the anatomical configuration of the synapse of these three BC classes. Two rod BC dendrites along with two flanking horizontal cell processes form a central invaginating profile in rod photoreceptor spherules. These are opposed to an electron dense ribbon with closely associated vesicles on the presynaptic side of the synapse. Similarly, a central cone DBC process flanked by two horizontal cell processes form invaginating profiles in cone pedicles, again opposed to presynaptic ribbons. In contrast, HBCs form flat contacts distal to the invaginating ribbon synapse on these same cone photoreceptors (see review by (Wassle et al., 2009)).

Mutations affecting rod photoreceptor presynaptic proteins that govern glutamate release generally disrupt the invaginating synapse, ribbon structure, and presumably glutamate release. These presynaptic disruptions are correlated with ectopic extensions of BC and HC dendrites that course through almost all of the ONL (Mansergh et al., 2005) (Chang et al., 2006) (Ball et al., 2002; Haeseleer et al., 2004; Wycisk et al., 2006). In contrast, most mutations affecting postsynaptic DBC proteins that govern postsynaptic signal transduction disrupt light signaling but do not alter the morphology of the invaginating ribbon synapse (Masu et al., 1995; Ball et al., 2003; Morgans et al., 2009; Koike et al., 2010; Peachey et al., 2012; Neuille et al., 2017). These DBC signaling complex components, hereafter referred to as the DBC signalplex, include mGluR6, TRPM1, GPR179, nyctalopin, RGS7, RGS11, R9AP, Gα0, Gβ13, Gβ5 and LRIT3.

Most studies have found that the dependence-hierarchy of DBC signalplex protein expression and functional interactions within the signalplex are similar in rod and cone DBCs. For example, TRPM1 expression depends on expression of nyctalopin (Pearring et al., 2011) and GPR179, the latter modulating the sensitivity of the DBC response (Ray et al., 2014). Based on the absence of TRPM1 in *Lrit3^nob6/nob6^* mouse retina, it was concluded that TRPM1 depends upon expression of LRIT3 (Neuille et al., 2015). Taken together, these two observations raise the question of whether LRIT3 or nyctalopin is the key protein required for TRPM1 trafficking to the dendritic tips of DBCs. To address this, we assessed expression of key signalplex proteins in an independently generated *Lrit3*^−/−^ mouse. Our results demonstrate that LRIT3 is required for nyctalopin expression and that mGluR6, TRPM1 and GPR179 are not required for LRIT3 expression. This absence of both nyctalopin and LRIT3 explains the no b-wave ERG phenotype in *Lrit3*^−^*^/^*^−^ mice.

Consistent with these data, we found that DBCs and ON RGCs in the ON signaling pathway of *Lrit3*^−^*^/^*^−^ retinas, lacked normal visual responses. Surprisingly, we found significant changes in the visual responses in the OFF pathway. OFF RGCs had reduced light-evoked responses, although response latency was similar to wildtype. The origin of these abnormally small responses is a reduced OFF BC light response. The OFF BC postsynaptic kainate receptors respond normally to exogenous application of agonist, which places the cause of OFF pathway signaling defects upstream, potentially due to altered photoreceptor glutamate release or clearance. LRIT3 is the first protein described whose absence impacts both ON and OFF BC function without apparent gross defects in synaptic architecture. These data suggest that LRIT3 has at least two functions: assembly of the postsynaptic DBC signalplex and control of the glutamate concentration in the synaptic cleft.

## MATERIALS AND METHODS

### Animals

All procedures were performed in accordance with the Society for Neuroscience policies on the use of animals in research and the University of Louisville Institutional Animal Care and Use Committee. Animals were housed in the University of Louisville AALAC approved facility under a 12 h/12 h light/dark cycle. The mouse line described in these studies, *Lrit3^emrgg1^*, is referred to as *Lrit3*^−^*^/^*^−^ throughout (see results for details). The phenotypes of all the other lines have been previously published. *Trpm1^−/−^, (Trpm1^tm1Lex^)*, (Shen et al., 2009); *Grm6^−/−^* (Masu et al., 1995); *Nyx*^nob^(Gregg et al., 2003); *GPR179^nob5^* (Peachey et al., 2012); *TgEYFP-Nyx* (Gregg et al., 2005); *MitoP-CFP* (Misgeld et al., 2007) *and TgVsx1-cerulean* (Hoon et al., 2015).

*Lrit3^+/+^* and *Lrit3^+/−^* litter mates of both sexes were used as controls throughout and the results from both were indistinguishable from C57Bl/6J. For all procedures, mice were anesthetized with a ketamine/xylazine solution (127/12 mg/kg, respectively) diluted in normal mouse Ringer’s or euthanized using CO**_2_** according to AVMA guidelines.

### Generation of *Lrit3*^−/−^ *mice with* Zinc Finger Nucleases (ZFN)

C3H/HeNTac/C57BL/6NTac hybrid embryos (363) were injected with 10 ng/μl *Lrit3* ZFN mRNA and 254 viable embryos were implanted into 9 Swiss Webster recipient mothers. Tail biopsies from offspring were collected and genomic DNA isolated using Direct Tail PCR solution (Thermo Scientific) supplemented with 0.2 μg/ml proteinase K (Thermo Scientific). Primers (5’-TAACCTGGGCATAGCCTGTC-3’; 5’-AAGGTCCAGGAAGGAGAAGG-3’) were used to amplify the ZFN targeted region (chr3:129503565, mm9). PCR fragments were either sequenced directly or cloned into the TopoBlunt vector (Invitrogen) and at least 10 clones sequenced. The *Lrit3*^−^*^/^*^−^ allele was backcrossed onto C57Bl/6J mice for 10 generations.

### Antibodies

Sheep anti-LRIT3 and rabbit anti-TRPM1 antibodies were generated by immunizing animals with peptides (LRIT3: AVTPSRSPDFPPRRII; TRPM1: SVVPEGQNTQQEKRSAETE) conjugated to KLH, by Biosynthesis Inc. (Lewisville, TX). Table 1 provides the details of all antibodies used, their dilutions and sources. The specificity of the LRIT3 and TRPM1 antibodies were validated by comparing immunostaining in control and *Lrit3***^−/−^** and *Trpm1***^−/−^** mice, respectively.

**Table 1.** Immunohistochemical reagents used in experiments

| Antigen | Dilution | Source | Catalogue # / reference |
| --- | --- | --- | --- |
| LRIT3 | 1:1000 | In house | Current paper |
| GPR179 | 1:2000 | In house | (Peachey et al., 2012) |
| TRPM1 | 1:1000 | In house | Current paper |
| Pikachurin | 1:2000 | Wako Chemicals, Richmond, VA | 011-22631;(Sato et al., 2008) |
| mGluR6 | 1:2000 | Gift, Dr. Kirill Martemyanov | (Cao et al., 2011) |

### Retina preparation for immunohistochemistry

Mice were killed by CO_2_ inhalation followed by cervical dislocation. Eyes were enucleated and the cornea and lens removed. The retina was dissected in PBS (pH 7.4) and fixed for 15–30 mins in PBS containing 4% paraformaldehyde, then washed in PBS for 5 mins, and cryoprotected in a graded series of sucrose solutions (5, 10, 15 and 20% in PBS) and finally in OCT:20% sucrose (2:1). The retinas were then frozen by immersion in an isopentane bath immersed in liquid nitrogen. Transverse 18 μm sections were cut on a cryostat (Leica Biosystems, Buffalo Grove, IL) and mounted on Superfrost Plus slides (Thermo Fisher Scientific, Waltham, MA). Slides were stored at −80°C until used in immunohistochemistry experiments.

Immunohistochemistry methods have been described previously (Hasan et al., 2016). Briefly, slides were dried at 37°C for 30 mins, rinsed in PBS for 5 mins and then in PBX (PBS + 0.5% Triton X-100) for 5 mins. Sections were then incubated in blocking buffer (PBX + 5% normal donkey serum) for 1 h followed by overnight incubation with primary antibody in blocking buffer. Sections were washed 3 × 10 mins in PBX, then incubated with secondary antibody diluted in blocking buffer for 1 h. Sections were washed 2 × 10 mins in PBX and 1 × 10 mins in PBS. Coverslips were mounted to slides using Vectashield (Vector Laboratories, Burlingame, CA). Sections were imaged on an FV-1000 Confocal Microscope (Olympus) with contrast and brightness adjusted using Fluoview Software (Olympus, Waltham, MA) or Photoshop (Adobe Systems, San Jose, CA).

### Electroretinography

ERG methods have been described previously (Ray et al., 2014). Briefly, mice were dark adapted overnight and anesthetized with a ketamine/xylazine solution (127/12 mg/kg, respectively) diluted in normal mouse Ringer’s and prepared for ERG recordings under dim red light. Pupils were dilated and accommodation relaxed with topical applications of 0.625% phenylephrine hydrochloride and 0.25% Tropicamide and the corneal surface anesthetized using 1% proparacaine HCl. Body temperature was maintained via an electric heating pad (TC1000 Temperature control, CWE Inc.). A clear acrylic contact lens with a gold electrode (LKC Technologies Inc.) was placed on the cornea and wet with artificial tears (Tears Again, OCuSOFT, Gaithersburg, MD). Ground and reference needle electrodes were placed in the tail and on the midline of the forehead, respectively. For scotopic responses, flashes (from −3.6 to 2.1 log cd s/m^2^) were presented to dark adapted animals. For photopic responses the animals were light adapted (20 cd/m^2^) for 5 mins and test flashes (from −0.8 to 1.9 log cd s/m2) were presented on this rod saturating background.

### Rod bipolar cell recordings

Methods for the preparation of retinal slices and whole cell recordings from bipolar cells have been described previously (Ray et al., 2014). Briefly, isolated retinas were placed on nitrocellulose paper (MilliporeSigma, Burlington MA). ~200 μm transverse slices prepared using a tissue slicer and placed in the recording chamber using vacuum grease. The recording chamber was constantly superfused with oxygenated Ringer’s and all solutions were maintained at 34–35°C. Recording electrodes with resistance measured between 6–9 MΩ were filled with Cs-gluconate intracellular solution (in mM: 20 CsCl, 107 CsOH, 107 D-Gluconic Acid, 10 NaHEPES, 10 BAPTA, 4 ATP, 1 GTP). 1% sulforhodamine was included in the intracellular solution to visualize and classify the cell based on its morphology (Ghosh et al., 2004). Inhibitory blockers (1 μM strychnine, 100 μM picrotoxin and 50 μM 6-tetrahydropyridin-4-yl methylphosphinic acid (TPMPA)) were included in bath solutions as was L-AP4 (4 μM) to saturate mGluR6 receptors. OFF cone BC somas were targeted for whole cell voltage clamp recording in Vsx1-cerulean reporter mice where Type 1 and 2 HBCs (BC1 and BC2) are sparsely labeled (Hoon et al., 2015). Only BC1s with an input resistance ~ 1 GΩ and access resistance < 25 MΩ were used for recording and were voltage clamped at +50 mV (Nawy, 2004; Shen et al., 2009). A Picospritzer II (Parker Instrumentation, Cleveland, OH) was used to pressure apply drugs onto BC dendritic tips located in the outer plexiform layer (OPL). Pressure applied drugs were the mGluR6 receptor antagonist α-cyclopropyl-4-phosphonophenylglycine (CPPG, 0.6 mM) to activate DBCs or kainate (50μm) to activate kainate receptors on HBCs. All reagents were purchased from Sigma-Aldrich, except L-AP4 and kainate, which were purchased from Tocris Bioscience (Avonmouth, Bristol, BS11 9QD United Kingdom). Clampfit 10.2 was used for off-line analyses of data. Currents were filtered off-line using a 20 Hz eight-pole Bessel low-pass filter.

### Glutamate imaging

To broadly target iGluSnFR expression to retinal ganglion cells and amacrine cells AAV2/1.*hSynapsin*.iGluSnFR in suspension was injected into the mouse eye, intravitreally (0.8 −1.0 μl volume H_2_O (0.8 − 3.0 × 10^13^ IU/μl). Animals were killed 14–21 d after AAV injection and the retinas prepared as described previously (Borghuis et al., 2011). Isolated retinas were mounted photoreceptor-side down on a nitrocellulose filter paper disc (Millipore Sigma, Burlington, MA) with 1.0-mm-diameter holes for visual stimulation through the condenser, and placed in a perfusion chamber on a custom-built two-photon fluorescence microscope. Tissue was continuously perfused with oxygenated Ames medium at physiological temperature (~6 ml/min; 34–36°C). Changes in iGluSnFR fluorescence, which represent the binding of glutamate to iGluSnFR, were measured as described previously (Borghuis et al., 2013), using a 60x, 1.0 NA, LUMPlanFl/IR objective (Olympus, Waltham, MA) and an ultrafast pulsed laser (Chameleon Ultra II; Coherent, Santa Clara, CA) tuned to 915 nm. The visual stimulus comprised a contrast reversing spot (150 μm diameter for BCs, 300 μm for GCs; 100% Michelson contrast; 1 Hz temporal modulation; 5 s duration) on a photopic background (λ_max_ = 395 nm; 5.8 × 10^4^ photons/μm^2/^s). Images (512 × 128 pixels) were acquired at 16 frames/sec; line scans were collected at 2 kHz and presented down-sampled to 0.5 kHz. Fluorescence responses were quantified using custom algorithms in Matlab (Mathworks, Natick, MA).

### RGC and OFF BC whole-cell recordings from retinal whole-mounts

Genetically identified BC1 cells were recorded in the whole mount retina of MitoP-CFP transgenic mice on a *Lrit3^−/−^* or control background. Alpha-type ON and alpha and delta-type OFF ganglion cells were targeted for recording based on soma size; cell type was verified post-hoc using two-photon fluorescence imaging of dye fills (Sulphorhodamine 101) and signature features of the recorded current responses. Visually-evoked excitatory and inhibitory currents were recorded in whole-cell configuration at the reversal potential for chloride (−69 mV) and cations (0 mV), respectively, using conventional methods (Multiclamp 700B, Digidata 1550, PClamp10; MDS Analytical Technologies, Union City, CA) and cesium-based internal pipette solution (in mM: 90 cesium methanesulfonate, 5 TEA-Cl, 10 HEPES, 10 BAPTA, 3 NaCl, 2 QX-314, 4 ATP-magnesium salt, 0.4 GTP-sodium salt, 10 Tris-phosphocreatine; pH 7.3, ~284 mOsm). OFF BC membrane voltage responses were recorded in current clamp, using potassium-based internal solution (in mM: 110 potassium methane sulfonate, 10 HEPES, 0.1 EGTA, 5 NaCl, 4 ATP-magnesium salt, 0.4 GTP-sodium salt, 10 Tris-phosphocreatine; pH 7.3, ~284 mOsm). Data was analyzed using custom algorithms in Matlab.

### Multi-electrode array recordings of RGCs

Methods of the recording procedures for the multi-electrode array (MEA) have been published (Peachey et al., 2017a). Mice were dark-adapted overnight, deeply anesthetized with ketamine/xylazine, and killed by cervical dislocation under dim red light. The retinas were dissected under dim red light in oxygenated Ringer’s solution. The vitreous was removed by incubation (10 min) in Ringer’s solution containing collagenase (12U/μl) and hyaluronidase (50U/μl) (Worthington Biochemicals, Lakewood, NJ). Pieces in dorsal and ventral retina were dissected (2 mm × 2 mm) and placed ganglion cell side down on a sixty electrode MEA (60MEA200/30Ti; Multi Channel Systems, Reutlingen, Germany). Retinal pieces were covered with a transparent cell culture membrane (ThermoFisher Scientific, Waltham, MA) and held in place with a platinum ring. The recording chamber was continuously perfused with oxygenated Ringer’s solution at 36°C throughout the experiment. Prior to recording, the preparation was allowed to settle for ~1hr in darkness. Spontaneous activity under dark adapted conditions was recorded for 10 mins followed by 10 or 20 trials of full-field light stimulation (5s/2s or 20s/10s interstimulus/stimulus interval, respectively). Light adapted responses were recorded after 5 min of adaptation to a background of 3.01 cd/m^2^. Three intensity levels were used and presented in order of increasing luminance (scotopic levels: 0.004, 0.03, 1.49 cd/m^2^ and photopic levels: 2.71, 14.7,303 cd/m^2^). Signals were band-pass filtered (80–3,000 Hz) and digitized at 25 kHz (MC Rack software; Multi Channel Systems, BW Germany). Spikes were recorded on individual electrodes. When electrodes recorded spikes from more than one RGC we sorted spikes using a principal components analysis (Offline Sorter; Plexon, Dallas, TX). Sorted units were exported, spikes binned (50 ms) and their spontaneous and visually evoked responses were analyzed (NeuroExplorer; Nex Technologies, Madison, AL). Spontaneous activity was examined for the presence of rhythmic activity using a power spectral density FFT analyses (NeuroExplorer) with 4096 frequency values and a Hann window function. The mean FFT was smoothed using a Gaussian filter with a bin width of 30 ms. The frequency of the FFT peak was plotted as a function of power in arbitrary units (A.U.). Using custom scripts, we defined light evoked responses as a response with a peak firing rate that was > +10 SEM above mean spontaneous. The time to peak of the evoked response was defined as the time after stimulus onset when the peak firing rate reached it maximus. Raster plots show responses to each stimulus presentations and peri-stimulus time histograms (PSTHs) represent the average spiking rate across all stimulus presentations.

### Statistical analyses

Prism 7.03 software (Graphpad Software, Inc., La Jolla, CA) was used to perform the statistical analyses as suited for the necessary comparison: two-way repeated measures ANOVAs, two-way ANOVAs, one-way ANOVAs, or t-tests. Tukey post-hoc tests were used when appropriate. Statistical significance = *P* < 0.05. When comparing Peak firing rate and time to peak data, Kruskal Wallis tests were used.

## RESULTS

We used zinc finger nucleases to generate a *Lrit3*^−^*^/^*^−^ mouse. Several lines were created and we used one with a 40 base pair deletion within exon 2 (Fig. 1A;(Ray, 2013) del chr3:129800675–129800714 GRCm38/mm10), which causes a frameshift. We assessed retinal function in the *Lrit3*^−^*^/^*^−^ mice with the electroretinogram (ERGs) and found that the a-wave was normal, but the b-wave was absent under both scotopic (Fig. 1B) and photopic (Fig. 1C) conditions. This no b-wave phenotype is similar to *Grm6*^−^*^/^*^−^ (Peachey et al., 2017), *Nyx^no^*^b^ (Pardue et al., 1998), *Trpm1*^−^*^/^*^−^ (Shen et al., 2009) and *GPR179^nob5/nob5^* (Peachey et al., 2012) mice. These results indicate that LRIT3 is critical to DBC function, similar to the results described for a different LRIT3 mutant mouse, *Lrit3^nob6/nob6^* (Neuille et al., 2014) and in humans, where mutations in LRIT3 cause complete congenital stationary night blindness (cCSNB)(Zeitz et al., 2013).

**Figure 1.**
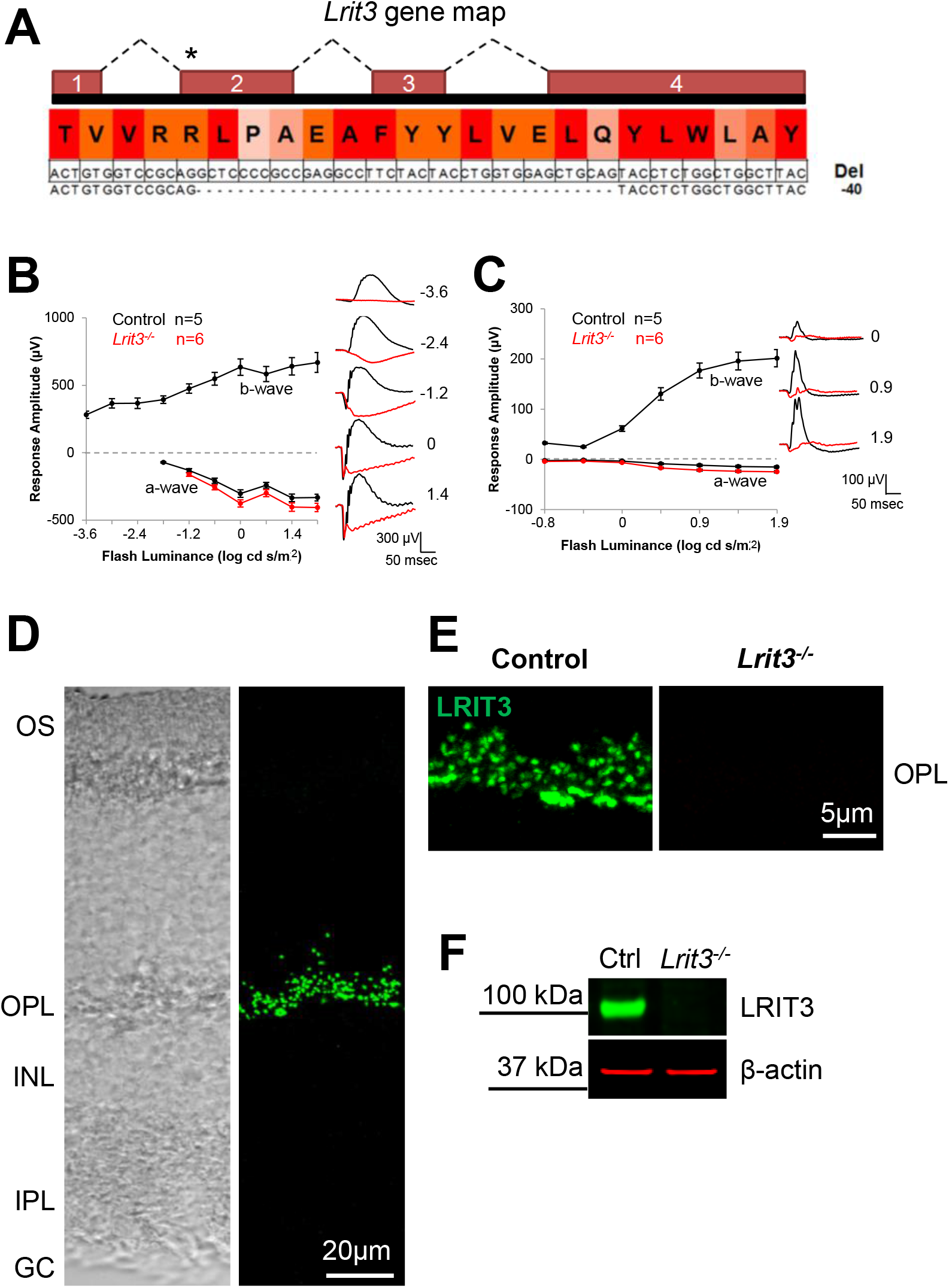
LRIT3 is required for normal ERGs and is expressed in the OPL. ***A***, Schematic of the *Lrit3* gene indicating the target region (*) for the zinc finger nuclease and resulting 40 bp deletion in the *Lrit3*^−^*^/^*^−^ mouse line. ***B***, Electroretinograms of control (black symbols) and *Lrit3*^−^*^/^*^−^ (red symbols) mice under scotopic and ***C***, photopic conditions. Example responses waveforms for 5 (scotopic) or 3 (photopic) luminance steps are shown, as well as summary data for all luminance steps. The *Lrit3*^−^*^/^*^−^ mice have a normal a-wave, but lack the b-wave under both scotopic and photopic conditions. ***D***, DIC (left) and immunohistochemical staining for LRIT3 in transverse sections from the control mouse retina. ***E***, Staining of OPL of control and *Lrit3*^−^*^/^*^−^ retinas for LRIT3. ***F***, Western blot for LRIT3 and a loading control β-actin in control and *Lrit3*^−^*^/^*^−^ retinas. These data show the LRIT3 antibody is specific. OS, outer segments, OPL, outer plexiform layer, INL, inner nuclear layer, IPL, inner plexiform layer, GC, ganglion cell layer.

To determine where the LRIT3 protein is expressed we generated an antibody to LRIT3 and verified its specificity in *Lrit3*^−^*^/^*^−^ retina using immunohistochemistry and western blotting (Fig. 1D-F). The LRIT3 antibody showed strong punctate staining exclusively in the OPL in the control retina and an absence of staining in the *Lrit3*^−^*^/^*^−^ retina (Fig. 1E). Western blot analyses further supported the absence of LRIT3 in retinal lysates from *Lrit3*^−^*^/^*^−^ retinas (Fig. 1F). These data show that the antibody is specific to LRIT3 and that expression of LRIT3 is restricted to the OPL.

### LRIT3 is required for expression of nyctalopin

Our previous work showed that the expression and correct localization of TRPM1 depends on the expression of nyctalopin (Pearring et al., 2011). The observation that TRPM1 is not expressed in *Lrit3*^−^*^/^*^−^ retina (Neuille et al., 2015) raised the question whether TRPM1 depends on LRIT3 directly or if its expression is lost due to a concomitant loss of nyctalopin. To address this, we examined the expression pattern of nyctalopin in *Lrit3*^−^*^/^*^−^ and control retinas (Fig 2A), by crossing a transgenic mouse line (*TgEYFP-Nyc*) that expresses an EYFP-nyctalopin fusion protein (Gregg et al., 2005) onto the *Lrit3*^−^*^/^*^−^ genetic background. In control *TgEYFP-Nyc* retina immunohistochemical analyses for EYFP-nyctalopin resulted in a punctate staining pattern in the OPL, localized to the tips of rod and cone DBCs, where it co-localized with both LRIT3 and the synaptic marker pikachurin (Fig 2Ai). *Lrit3*^−^*^/^*^−^*/TgEYFP-Nyc* retina showed no EYFP-nyctalopin expression (Fig. 2Aii), even though these mice carried a copy of the EYFP-nyctalopin transgene. Thus, we conclude that nyctalopin expression on the DBC dendritic tips depends on LRIT3 expression.

**Figure 2.**
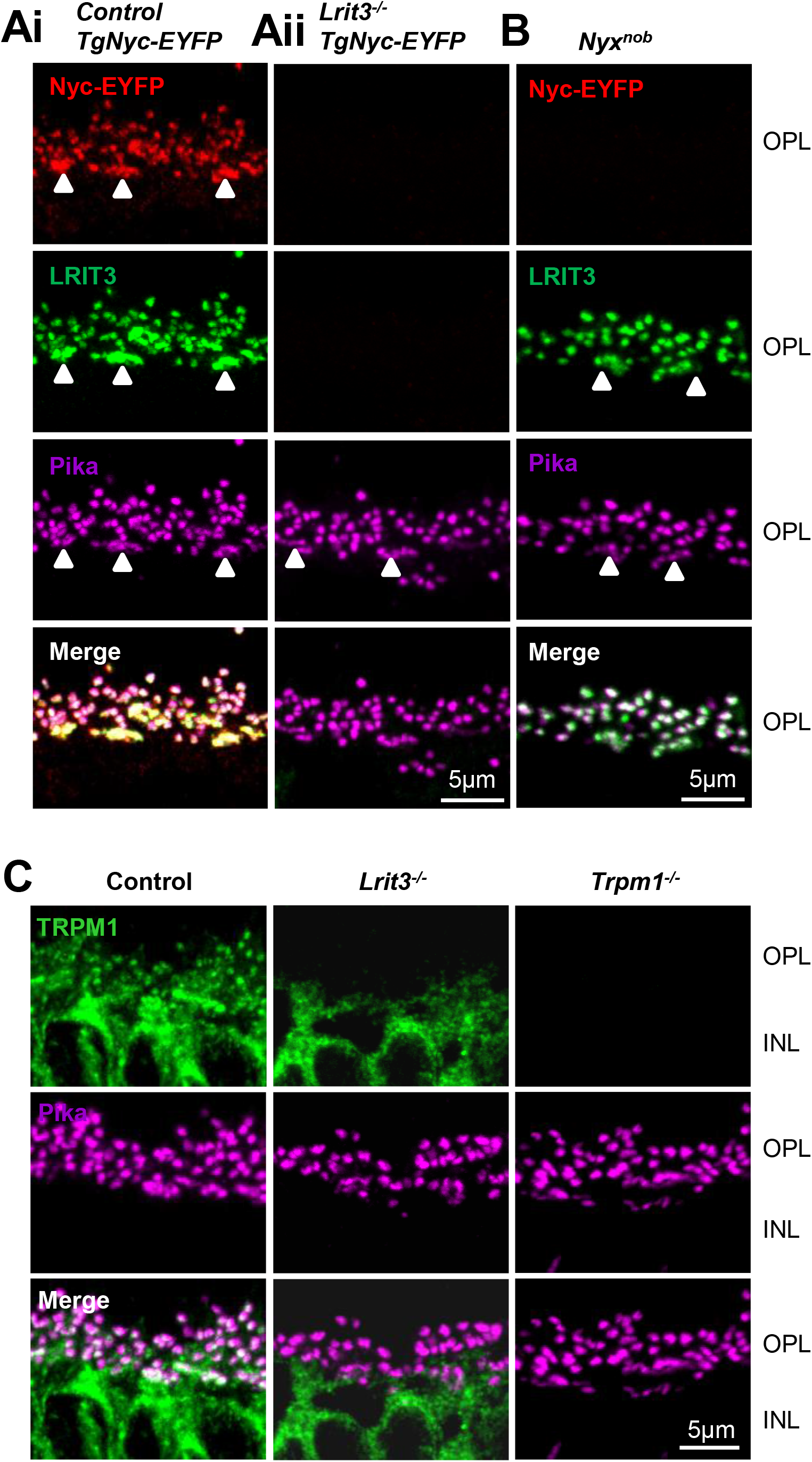
Nyctalopin expression requires LRIT3 expression. ***A***, EYFP-Nyctalopin (red), LRIT3 (green) and pikachurin (magenta) staining in ***Ai***, control and ***Aii****, Lrit3*^−^*^/^*^−^ retinas, carrying the TgNyc-EYFP transgene. Note that Nyctalopin-EYFP is not expressed in the *Lrit3*^−^*^/^*^−^ retina. ***B***, Although *Nyx^nob^* retinas lack nyctalopin, LRIT3 is expressed and localized to the dendrites of the rod and cone (marked by arrowheads) DBCs. ***C***, As expected, TRPM1 (green) is mislocalized in *Lrit3*^−^*^/^*^−^ retinas and pikachurin expression is normal. These are representative images of data from at least 4 mice. OPL, outer plexiform layer; INL, inner nuclear layer.

We tested whether LRIT3 expression depends on nyctalopin by examining its expression in *Nyx^nob^* mutant mice. In *Nyx^nob^* retina (Fig. 2B), both LRIT3 and pikachurin are expressed and co-localize on the rod and cone DBC tips, similar to control mice (Fig 1Ai). We also verified the absence of TRPM1 expression in our independent line of *Lrit3*^−^*^/^*^−^ mice (Fig 2C).

Based on these observations we conclude that nyctalopin expression depends on LRIT3 expression, and that the loss of TRPM1 from the dendritic tips of all *Lrit3*^−^*^/^*^−^ DBCs (Fig. 1E) results from the loss of nyctalopin expression.

### LRIT3 localization is independent of expression of other DBC cascade components

The absence of mGluR6, GPR179 or TRPM1 has varying effects on the localization and expression of other cascade components (see (Gregg et al., 2014) for review). We determined whether these DBC signalplex components are required for normal expression of LRIT3, using *Grm6*^−^*^/^*^−^, *Gpr179*^−^*^/^*^−^ and *Trpm1*^−^*^/^*^−^ mouse retinas. Sections from each line were reacted with antibodies to pikachurin and LRIT3, and the staining patterns compared to control and *Lrit3*^−^*^/^*^−^ retinas (Fig. 3). Both LRIT3 and pikachurin are localized normally to the dendritic tips of both rod and cone DBCs in *Grm6*^−^*^/^*^−^*, Gpr179*^−^*^/^*^−^ and *Trpm1*^−^*^/^*^−^ mouse retina. These data demonstrate that the trafficking and localization of LRIT3 is independent of mGluR6, GPR179 and TRPM1.

**Figure 3.**
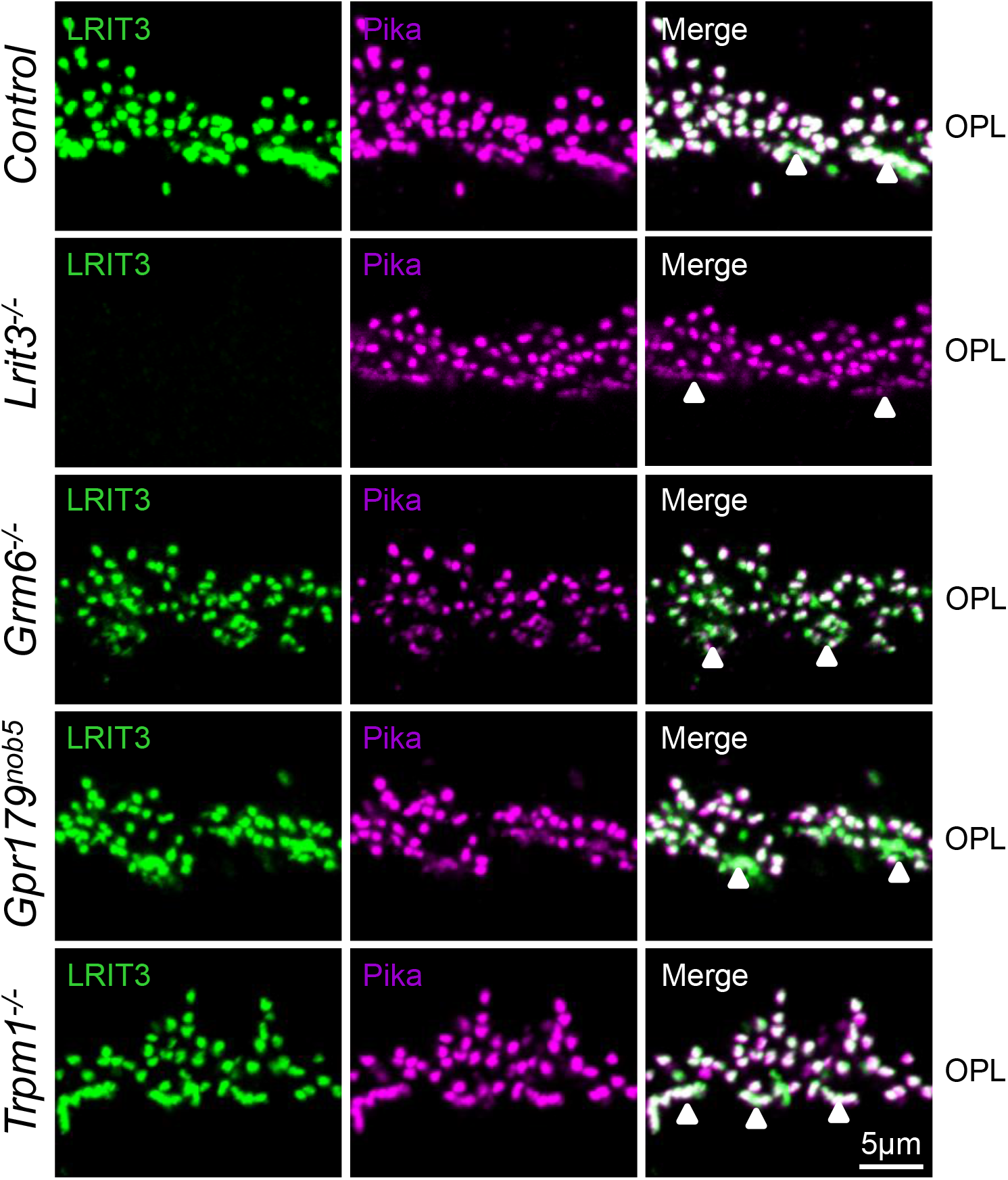
LRIT3 and pikachurin are expressed normally in the OPL of *Grm6*^−^*^/^*^−^*, Gpr179^nob5^* and *Trpm1*^−^*^/^*^−^ mouse retinas. Retinas from control, *Lrit3*^−^*^/^*^−^*, Grm6*^−^*^/^*^−^*, Gpr179^nob5^* and *Trpm1*^−^*^/^*^−^ mouse retinas were immuno-stained for LRIT3 (green) and pikachurin (magenta). The merged images show that LRIT3 and pikachurin co-localize. Arrowheads indicate cone terminals. OPL, outer plexiform layer.

The *Lrit3^nob6^* mutant is reported to lack expression of TRPM1 in rod BC dendrites and TRPM1, mGluR6, GPR179, RGS7 and 11, Gβ5 and GαO in cone BC terminals (Neuille et al., 2015). We found similar results in our *Lrit3*^−^*^/^*^−^ line and extend the previous report with R9AP, which also is missing at the cone, but not rod BC terminals (Fig. 2 and Extended Fig. 2.1).

### RGC signaling in the *Lrit3***^−^*^/^*^−^** retina is abnormal

We used the same assays of RGC function that defined abnormal response properties of RGCs in other (no b-wave) mouse models of cCSNB, *Nyx^nob^*, and *Grm6*^−^*^/^*^−^ (Demas et al., 2006) (Peachey et al., 2017a). These abnormalities include the absence of normal latency ON responses (light onset) and rhythmic bursting in almost all RGCs. However, OFF responses had normal latency.

We surveyed the response properties of >1000 RGCs in *Lrit3*^−^*^/^*^−^ and control retinas using a multi-electrode array (MEA). To optimize stimulus conditions we evaluated responses in a subset of RGCs over 3 intensity flashes in the dark-adapted and light adapted retina. In control retinas, the brightest stimulus intensity (303 cd/m^2^) evoked responses in ~100% of RGCs (Fig. 4A). Conversely, only 39% of *Lrit3*^−^*^/^*^−^ RGCs produced a light response, even at the brightest stimulus intensity. For the remaining experiments we present data generated using the brightest photopic flash stimulus. We divided RGCs into (1) non-responsive RGCs, which showed spontaneous activity but no visually evoked response, and among visually responsive RGCs, (2) those with a response to light onset (ON or delayed ON (dON)), (3) those with a response to light offset (OFF) or (4) those with a response to light onset and offset (ON/OFF). We compared the distribution of these four type in control, *Lrit3*^−^*^/^*^−^, and *Grm6*^−^*^/^*^−^ retinas (summarized in Fig. 4B; representative visually evoked responses for each genotype shown in Fig. 4C). All control RGCs had excitatory light-evoked spiking responses significantly above their spontaneous activity. Of these control RGCs, 47% had an excitatory response at light onset (ON RGCs), 27% had an excitatory response at light offset (OFF RGCs), and 20% had an excitatory response to both luminance changes (ON/OFF RGCs).

**Figure 4.**
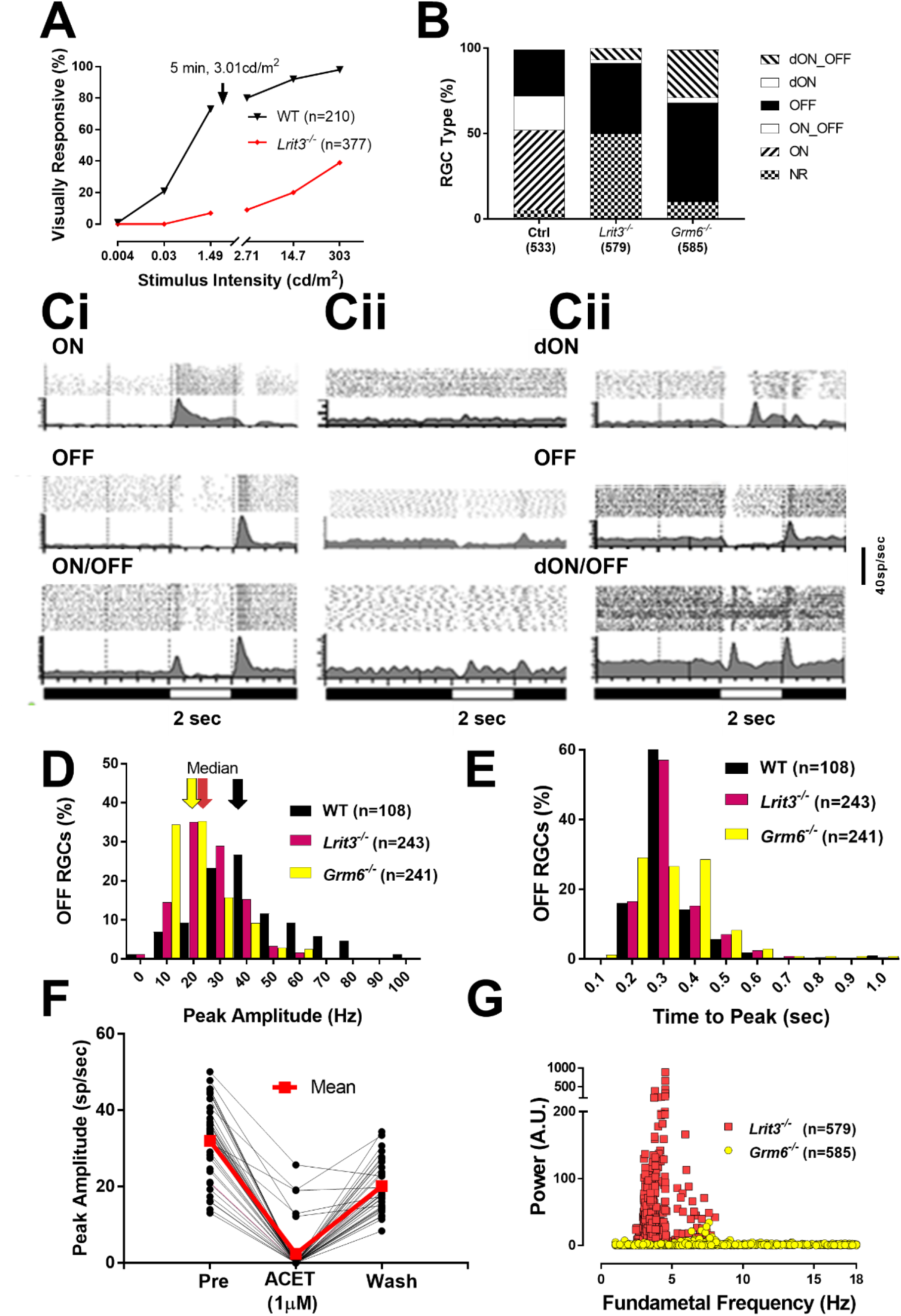
The visual responses of *Lrit3*^−^*^/^*^−^ RGCs are significantly altered compared to controls and *Grm6*^−^*^/^*^−^. ***A***, Intensity response function of RGCs in control and *Lrit3*^−^*^/^*^−^ RGCs. The break in the x-axis represents a 5 min light adaptation to 3.01 cd/m^2^. ***B***, RGC functional classes in control, *Lrit3*^−^*^/^*^−^ and *Grm6*^−^*^/^*^−^ retinas. The total number of RGCs for each genotype is shown and the data were obtained from 12 retinal pieces from 8 control mice; 16 retinal pieces from 6 *Lrit3*^−^*^/^*^−^ mice; and 10 retinal pieces from 4 *Grm6*^−^*^/^*^−^ mice. There are significantly more (50%) visually non-responsive; defined as cells with spontaneous but no visually evoked activity, RGCs in the *Lrti3*^−^*^/^*^−^ mice than either control or *Grm6*^−^*^/^*^−^. (Fishers exact test, p<0.001 for both comparisons). ***C***, Representative average peristimulus time histograms (above - raster plots to individual stimulus presentation) of responses recorded on a multielectrode array to a full field light stimulus (303 cd/m^2^) on a 0.3 cd/m^2^ background. ***Ci***, Control RGC responses can be classified as ON, OFF and ON/OFF based on whether spiking peaks to light onset, offset or both, respectively. All responses occur < 0.4 sec after stimulus onset. ***Cii*** and ***Ciii****, Lrit3*^−^*^/^*^−^ and *Grm6*^−^*^/^*^−^ RGC responses can be classified into the same general groups but the time to the peak response to light onset is > 0.4 sec and these responses are referred to as delayed ON (dON). Among visually responsive RGCs dON RGCs are only found in *Lrti3^−/−^* and *Grm6*^−^*^/^*^−^ mutant retinas. ***D***, The peak responses of OFF RGCs are decreased in *Lrit3*^−^*^/^*^−^ and *Grm6*^−^*^/^*^−^ compared to controls (median: 25 and 19 Hz, respectively; compared to 38Hz in controls, Kruskal-Wallis followed by Dunn’s test; p < 0.001 for both comparisons). ***E***, Distribution of response latencies in the OFF RGCs in *Lrit3*^−^*^/^*^−^ and *Grm6*^−^*^/^*^−^ is not different than control (Kruskal-Wallis p=0.995). ***F***, OFF responses in RGCs are mediated via kainate receptors. Responses show 37 RGCs peak response (sp/sec), before (32±1.6 SEM)), during (2.4±1.1) and after ACET (20.11±1.1) treatment. ***G***, Fast Fourier transform to determine rhythmicity of the spontaneous activity of RGCs from *Lrit3*^−^*^/^*^−^ and *Grm6*^−^*^/^*^−^ RGCs. Data shows many *Lrit3*^−^*^/^*^−^ cells exhibit rhythmic firing, whereas few do so in the *Grm6*^−^*^/^*^−^ retinas.

Against this baseline we compared RGC responses in *Lrit3*^−^*^/^*^−^ mice, and also in *Grm6*^−^*^/^*^−^ mice because they share the no b-wave phenotype. The percentages of RGC response types in the two mutants were strikingly different and each also differed from control (Fig 4B). As expected from a no b-wave phenotype, neither *Lrit3*^−^*^/^*^−^ nor *Grm6*^−^*^/^*^−^ retinas had ON responses with short latency, although each had RGCs whose ON responses were significantly delayed compared to control (> 0.5 sec, time to peak), 9% in *Lrit3*^−^*^/^*^−^ retinas compared to 31% in the *Grm6*^−^*^/^*^−^ retinas. We also found a large and unexpected difference in the percentage of visually non-responsive RGCs in the *Lrit3*^−^*^/^*^−^ versus *Grm6*^−^*^/^*^−^ retinas (50% vs. 10%, Fishers p<0.0001).These differences in the proportions of non-responsive RGCs as well as across visually responsive classes prompted us to examine additional aspects of their responses.

The large number of NR cells in the *Lrit3*^−^*^/^*^−^ suggests that their sensitivity may be impacted, so we examined the peak response of RGCs and found that *Lrit3*^−^*^/^*^−^ OFF RGC peak responses were significantly smaller than control (median = 24.7sp/sec, n=242 vs. 37.6 sp/sec, n=86) although *Lrit3*^−^*^/^*^−^ OFF responses were larger than *Grm6*^−^*^/^*^−^ RGCs (18.5 sp/sec, n=241; Fig 4D; Kruskal-Wallis; both comparisons p < 0.0001). In contrast, the time to peak response of OFF RGCs was similar across genotype (Fig. 4E; Kruskal-Wallis; p = 0.995). In the few *Lrit3*^−^*^/^*^−^ ON/OFF RGCs the time to peak firing at light onset (median = 660 msec, n=8) was significantly delayed compared to control (median = 250 ms, n=263; p=?) and similar to *Grm6*^−^*^/^*^−^ RGCs (median = 550 ms, n=35; p=?) we reported previously (Peachey et al., 2017b).

To verify that the OFF RGCs responses in *Lrit3*^−^*^/^*^−^ were initiated in the OFF pathway we compared light evoked responses in control solution to those in the presence of the kainate receptor antagonist ACET (1 μM), and after 1 hour wash (Fig 4F). Of the 515 *Lrit3*^−^*^/^*^−^ RGCs, 180 (35%) had visually evoked OFF responses in control solution. After addition of ACET to the bath, only 11 (6%) retained a visual response, and all showed a reduced peak amplitude (~43% of control). After washout, 37 RGCs recovered their OFF response. These data demonstrate that light-evoked OFF responses originated in the OFF signaling pathway.

We previously reported that *Nyx^nob^* RGCs show rhythmic bursting in their spontaneous activity, which also is superimposed on their light evoked responses (Demas et al., 2006). Both the post stimulus time histograms, as well as the raster plots (Fig 4Cii) show that rhythmic activity is present in *Lrit3*^−^*^/^*^−^ RGCs. *Lrit3*^−^*^/^*^−^ RGCs showed rhythmic activity regardless of whether a visual response was present or absent. Similar to *Nyx^nob^* RGCs, *Lrit3*^−^*^/^*^−^ RGCs showed rhythmic activity with a fundamental frequency between 3 and 8 Hz (Fig 4G). In contrast, the majority of *Grm6*^−^*^/^*^−^ RGCs did not have rhythmic activity and in the few that did (Fig. 4G, 24/866), the fundamental frequency of the response modulation ranged between 5.6 and 9 Hz and the modulation amplitude was lower by ~3-fold compared with *Lrit3*^−^*^/^*^−^ RGCs (11.4 vs 34.8 A.U).

### ON and OFF BC signaling in the *Lrit3***^−^*^/^*^−^** retina is abnormal

The large number of non-responsive *Lrit3*^−^*^/^*^−^ RGCs and OFF RGCs with reduced peak firing rates led us to examine light-evoked glutamate release in the IPL at the level of the bipolar cell axon terminals. We expressed the fluorescent glutamate sensor iGluSnFR in RGCs and amacrine cells using viral transduction and recorded iGluSnFR fluorescence during visual stimulation, using two-photon imaging (Borghuis et al., 2013). Control retinas showed robust fluorescence responses in the ON and OFF sublaminae of the IPL, reflecting glutamate release from ON and OFF BCs, respectively (Fig 5A, B). In *Lrit3*^−^*^/^*^−^ retina glutamate release was negligible in the ON sublaminae (*Lrit3*^−^*^/^*^−^ −0.002 ± 0.009, n = 8 *vs*. control 0.306 ± 0.028, n = 9; t-test, p<0.0001), as expected from the ERG no-b wave phenotype. Although we observed clear stimulus modulated glutamate release in the OFF sublaminae at light decrements, the amplitude of the response (ΔF/F) was reduced compared with control (*Lrit3*^−^*^/^*^−^, 0.173 ± 0.037, n = 11 *vs*. control 0.420 ± 0.065, n = 5; t-test, p = 0.0027; Fig. 5B, C).

**Figure 5.**
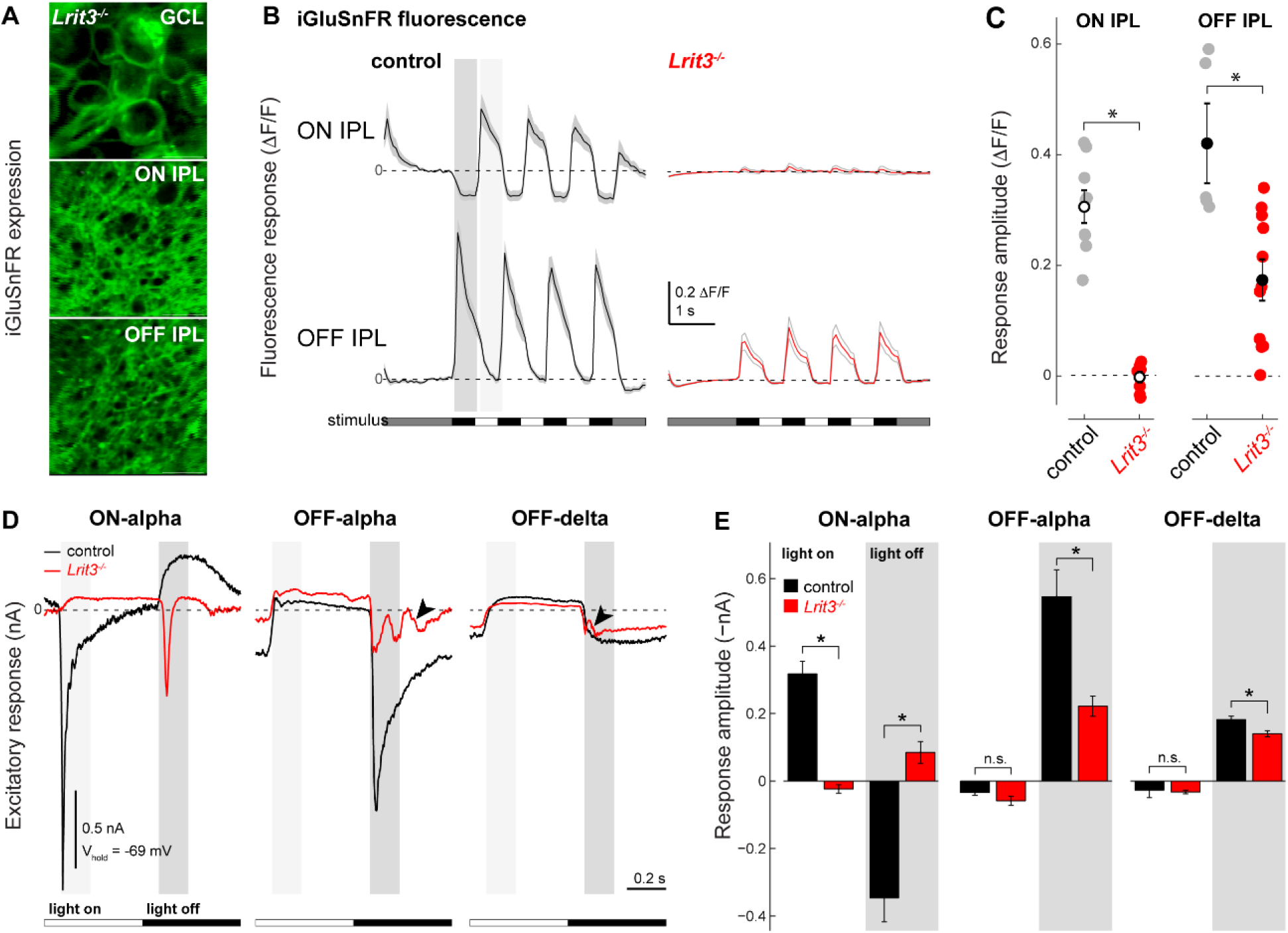
Three types of *Lrit3*^−^*^/^*^−^ RGCs show decreased excitatory input to light stimuli. ***A***, iGluSnFR expression in the GC, and ON and OFF sublaminae of the IPL following viral transduction with AAV2/1-*hsyn*-iGluSnFR. ***B***, Visually-evoked changes in glutamate levels as detected by iGluSnFR fluorescence in the ON (top traces) and OFF (bottom traces) layers of the IPL for control (black traces) and *Lrit3*^−^*^/^*^−^ (red traces), to a contrast reversing spot on a photopic background (1.2 × 10^4^ R/rod and cone/s; 150 μm diameter, 1Hz square wave, 100% Michelson contrast). ***C***. Quantification of the change in iGluSnFR fluorescence in the ON and OFF sublaminae of control and *Lrti3*^−^*^/^*^−^ retinas for all recorded areas (ON, control n = 9, *Lrit3*^−^*^/^*^−^ n = 8; OFF, control n = 5, *Lrit3*^−^*^/^*^−^ n = 11). ***D***, Electrophysiological whole-cell recordings of light-evoked current responses of three identified ganglion cell types (same stimulus as in *B*, except diameter 350 μm; recorded in voltage clamp mode at the reversal potential for chloride, −69 mV). ***E***, Quantification of the light-evoked current response amplitude for all recorded cells. ON and OFF response amplitudes were computed as the mean of the recorded current following a light increment and decrement, respectively (light and dark gray regions in *D*). For all responses the amplitude of the excitatory (inward) current is inverted for ease of interpretation. Data were compared using a t-test, * p< 0.05.

To understand the changes underlying the reduced signaling in ON and OFF pathways at the level of the inner retina, we examined three well characterized RGCs—ON alpha, OFF alpha and OFF delta— using whole-cell voltage clamp recordings in control and *Lrit3*^−^*^/^*^−^ retinal whole mount preparations (Fig. 5D). RGC identity was confirmed based on morphology defined by a red fluorescent dye that diffused from the intracellular pipette solution. Clamped at the reversal potential for chloride (E_CI_, −69 mV), control ON alpha RGCs showed a robust inward (excitatory) current at light onset and suppression of the tonic inward current at light offset, reflecting the stimulus-evoked modulation in glutamate release from presynaptic ON BCs. In *Lrit3*^−^*^/^*^−^ ON alpha cells, both the inward ON response and the suppression of the tonic inward current were absent (inward current *Lrit3*^−^*^/^*^−^ 23.5 ± 12.3 pA, n = 14 *vs*. control −317.1 ± 37.6 pA, n = 6; t-test, p<0.0001; Fig. 5E). Instead, the suppression of glutamate release following light offset was replaced by a small transient inward current, which is likely due to AII amacrine-cell mediated signaling from OFF BCs to ON BCs, but was not investigated further. OFF alpha and OFF delta RGCs in control retinas each showed their characteristic inward currents at light offset (Borghuis et al., 2014). While the response polarity and timing in *Lrit3*^−^*^/^*^−^ OFF alpha and delta cells were similar to control, the response amplitudes of both cell types was significantly decreased (OFF alpha: *Lrit3*^−^*^/^*^−^ − 221.4 ± 29.6 pA, n = 7 *vs*. control −546.6 ± 78.8 pA, n = 6; t-test, p = 0.0017; OFF delta −139.7 ± 9.2 pA, n = 3 vs. −181.8 ± 10.4 pA, n = 3; Fig. 5E). The observation that excitatory input to the transient OFF cell decreases 60%, whereas the decrease in the sustained OFF cell is just 23% may indicate that loss of LRIT3 differentially affects the specific BC types presynaptic to each RGC type.

These data show that *Lrit3*^−^*^/^*^−^ ON alpha RGCs lack excitatory synaptic input at light onset, as expected, but also that the amplitudes of light-evoked excitatory current in at least two identified OFF RGC types are reduced. Collectively, the MEA and voltage clamp data suggest that all RGCs in *Lrit3*^−^*^/^*^−^ retinas are less responsive than controls, although the magnitude of the impact varies with RGC type. Given that LRIT3 is not expressed in the IPL (Fig. 1D), the most likely explanation for the defects in OFF RGCs is that their OFF BC input is decreased.

### *Lrit3*^−^*^/^*^−^ retinas show abnormal ON and OFF BC function

We used patch clamp recordings of rod BCs and measured TRPM1 mediated currents. We simulated dark by bathing the slices in L-APB to activate mGluR6 and close TRPM1, then inactivated the mGluR6 to TRPM1 cascade by puffing on CPPG, which opens TRPM1 (Shen et al., 2009). In control cells CPPG mediates a robust current that is absent in both *Lrit3*^−^*^/^*^−^ and *Trpm1*^−^*^/^*^−^ rod BCs (Fig. 6A). The small residual current in both knockouts has been observed previously and arises from an unknown source (Ray et al., 2014). The decreased amplitude of the OFF RGCs in the MEA and patch clamp recordings of *Lrit3*^−^*^/^*^−^ RGCs, and the decrease in glutamate release in their OFF sublaminae could result from disruption of the AMPA/kainate receptors at the cone:cone DBC synapse. Thus, we recorded responses to kainate puffs in Type 1 OFF BCs (BC1s) in retinal slices from *Tg-Vsx/Lrit3*^−^*^/^*^−^ retinas (Fig. 6B). Kainate elicited a robust inward current in control BC1s. On average the control response was similar to the kainate response in *Lrit3*^−^*^/^*^−^ BC1s. Our interpretation is that the absence of LRIT3 does not impact either the number or function of kainate receptors on BC1s. In addition, we examined light-evoked currents in identified BC1s using two-photon fluorescence imaging in a whole mount retina preparations. Whole-cell recordings in response to light stimuli at a range of holding potentials, showed robust light-evoked excitatory and inhibitory currents (Fig 6C). Responses were characterized by an excitatory current following light decrements recorded at E_CI_ (~-60 mV) and an inhibitory current at E_cation_ (~0 mV) following both light decrements and increments, indicating that BC1s receive a combination of feed-forward (OFF pathway-mediated) and cross-over inhibition (ON pathway-mediated through the AII amacrine cell). In *Lrit3*^−^*^/^*^−^ BC1s, the excitatory currents were decreased dramatically and the transient part of the response was absent (Fig. 6D,E). Since the iGluR density appears normal (Fig. 5B), this results suggests that either glutamate release is decreased or its concentration changed in the cleft. Finally, *Lrit3*^−^*^/^*^−^ BC1 inhibitory currents were less synchronized to the stimulus, consistent with loss of function in the ON signaling pathway responsible for cross-over inhibition.

**Figure 6.**
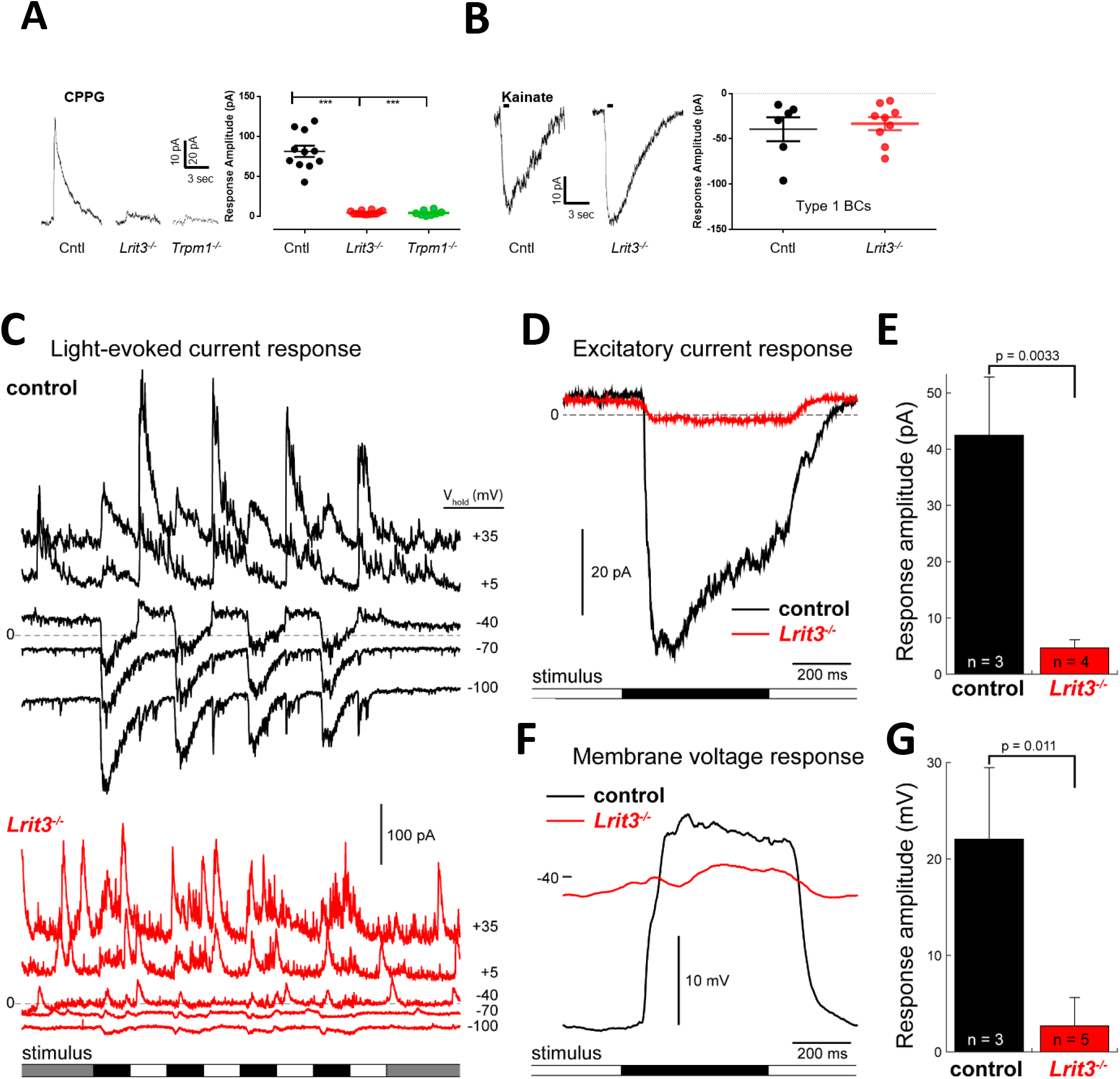
*Lrit3*^−^*^/^*^−^ rod BCs lack functional TRPM1 responses and BC1 cells have decreased excitatory input. ***A***, Patch clamp recordings from rod BC from control, *Lrit3*^−^*^/^*^−^ and *Trpm1*^−^*^/^*^−^ mice in response to puffs of the mGluR6 antagonist CPPG (3mM). Summary data from multiple cells shows a dramatically decreased response in *Lrit3*^−^*^/^*^−^ and *Trpm1*^−^*^/^*^−^ cells, consistent with absence of the ERG b-wave in these knockouts. (*** ANOVA, F(2,30),p < 0.001, Tukey post-hoc tests, P< 0.001 for control vs both knockouts). ***B***, Responses of BC1 cells to kainate puffs (50μm) in retinal slices from control and *Lrit3*^−^*^/^*^−^ retinas. The summary data shows there is no difference in the maximal responses between the two genotypes (t-test, p=0.67). ***C***, Electrophysiological whole-cell recordings of a genetically identified BC1 cells in control (top) and *Lrit3*^−^*^/^*^−^ (bottom) mouse whole-mount retina. Traces represent the current recorded in voltage clamp mode at different holding potentials (E_CI_ = −69 mV; E_cat_ = 0 mV). *Lrit3*^−^*^/^*^−^ showed a marked absence of excitatory current (lower traces). ***D***, Light-evoked excitatory current response of an example control and *Lrit3*^−^*^/^*^−^ BC1. Traces show the average response to nine repeats of the square wave, contrast-reversing spot stimulus presented on a photopic background (1.2 · 10^4^ R^*^//rod and cone//s; 150 μm diameter, 1Hz square wave, 100% Michelson contrast). ***E***, Quantification of the excitatory current response for all recorded BC1 cells (data partially shown in *D*). ***F, G***, As D and E, for membrane voltage response recorded in current clamp mode. Statistics in ***E,G***, t-test.

Membrane voltage responses recorded in control BC1s in current clamp mode showed a ~20mV membrane depolarization at light offset, which was only 1–2mV in *Lrit3*^−^*^/^*^−^ BC1s. This demonstrates that the absence of LRIT3 causes near-complete loss of function in this bipolar cell type. Because BC1s are the predominant source of excitatory input to OFF delta GCs, this result explains their loss of excitation (Fig. 5). Since the OFF alpha cells sample from a different complement of BC types (primarily BC3a, BC3b, and BC4) their decreased excitatory input (Fig. 5B, C) suggests that loss of function demonstrated in BC1 generalizes to other OFF BC types, including those that drive OFF alpha GCs. The decreased response of all OFF BCs is consistent with the decrease in glutamate release in the OFF sublaminae (Fig 5B). We propose that LRIT3 is critical for normal glutamate concentration dynamics within the cone➔cone BC synaptic cleft.

## DISCUSSION

We show that LRIT3 is required not only for ON but also, surprisingly, for OFF BC function, independent of crossover input from the ON pathway. The latter defect leads to abnormal OFF RGC responses that have not been observed in any other CSNB1 mouse model examined to date. Mutations in *LRIT3* were first identified after exome sequencing of DNA samples from individuals with CSNB1 (Zeitz et al., 2013). Our ERG studies, and those of others (Fig. 1, (Ray, 2013; Neuille et al., 2014)) show the typical no b-wave phenotype reported in numerous other CSNB1 mouse models (Masu et al., 1995; Pardue et al., 1998; Morgans et al., 2009; Shen et al., 2009; Koike et al., 2010; Peachey et al., 2012).

LRIT3 colocalizes with DBC signalplex proteins: TRPM1, mGluR6, GPR179, nyctalopin, RGS7, RGS11, Gβ5 and R9AP. Analyses of *Lrit3*^−^*^/^*^−^ and *Lrit3^nob6/nob6^* mice show that LRIT3 is required for localization of TRPM1 to the rod DBC signalplex. On cone DBCs LRIT3 also is required for TRPM1 localization, but also all the other signalplex proteins (Fig. 2 and 2.1, (Ray, 2013) (Neuille et al., 2015). Based on this result, Neuille and colleagues (Neuille et al., 2015) argued that LRIT3 was required for TRPM1 localization. Our data show that this is true, but that the hierarchy must include nyctalopin expression, which is directly required for TRPM1 trafficking/localization (Pearring et al., 2011). In this scenario, the loss of TRPM1 is secondary to the loss of nyctalopin in the *Lrit3*^−^*^/^*^−^ retina.

Further, PNA staining, which marks cone terminals in the OPL, also is absent or greatly reduced in expression. These data indicate LRIT3 has additional functions at the cone to cone DBC synapse, compared to rod to rod bipolar cells. Since PNA is thought to be a presynaptic marker, LRIT3 is the first protein whose absence not only causes DBC dysfunction but also alters expression of a presynaptic protein, without structural disruption of the synapses.

This disruption in PNA expression, led us to examine dysfunction in OFF BC and RGC function in *Lrit3*^−^ mice. RGC response properties have been examined in several CSNB1 mouse lines *(Nyx^nob^ (Demas et al., 2006), Grm6*^−^*^/^*^−^ *(Renteria et al., 2006; Pinto et al., 2007; Peachey et al., 2017a), Trpm1*^−^*^/^*^−^ (Takeuchi et al., 2018) *and Lrit3*^−^*^/^*^−^ (Fig. 4) and *Lrit3^nob6^* (Neuille et al., 2017). Consistent with the absence of an ERG b-wave these mutant RGCs lack normal ON responses (MEA and patch clamp recordings). The only responses to light onset are significantly delayed and have been shown to arise from the OFF BCs, via crossover pathways (Renteria et al., 2006). In the *Lrit3*^−^*^/^*^−^ retinas we find that the response amplitude of the OFF RGCs is decreased compared to controls and that over half the RGCs fail to respond to light stimuli. These results differ from a previous report using the *Lrit3^nob6/nob6^* mouse line (Neuille et al., 2017).

The absence of ON BC crossover input to RGCs results in decreased response amplitudes in both *Lrit3*^−^*^/^*^−^ and Grm6^−^*^/^*^−^ retinas (Fig. 4), and similar effects can be mimicked by addition of L-AP4 to control retinas. However, a unique feature of *Lrit3*^−^*^/^*^−^ RGCs is that there are 5 times as many visually non-responsive RGCs compared to *Grm6*^−^*^/^*^−^ retinas (50% vs 10%). This suggests that the underlying mechanisms responsible for the decreased responses differ. The recordings of excitatory currents in two types of OFF RGCs, OFF-alpha and OFF delta, support this idea; both show significant decreases in light responses in the *Lrit3*^−^*^/^*^−^ compared to control. Of particular note is that the decrease in *Lrit3*^−^*^/^*^−^ OFF-alpha cells (~90%) is far greater than the decrease produced when WT OFF-alpha cells are recorded in the presence of L-AP4 (~25% decrease in excitatory current (Borghuis et al., 2014)). Further, when OFF delta cells are recorded in the presence of L-AP4 their excitatory currents actually increase (Borghuis et al., 2014). From these data we postulated that the effects on the *Lrit3*^−^*^/^*^−^ OFF RGCs results from a direct decrease in excitatory input from OFF BCs to OFF RGCs. This is fully supported by our recordings from one type of OFF BC, BC1’s, in which the light responses are significantly decreased in the *Lrit3*^−^*^/^*^−^ compared to control retinas.

The mechanism underlying the decreased excitatory input to OFF BCs requires further study. That said, it is clear that it does not arise from defects in photoreceptor transduction because the *Lrit3*^−^*^/^*^−^ scotopic and photopic a-waves are the same as controls (Fig. 1). Because the expression of LRIT3 is restricted to the OPL, we propose two possible changes that result in reduced excitatory responses in OFF BCs. First, LRIT3 could modulate photoreceptor glutamate release, although at the rod synapse this requires a mechanism independent of the calcium channel Cav1.4. Neuille et. al (2017) reported that the number of invaginating cone DBCs was decreased without effecting the number of flat synapses with OFF BCs in the *Lrit3^nob6/nob6^* retina. Therefore, it is possible a disruption in the synaptic architecture could negatively impact glutamate release. The other possibility is that LRIT3 is involved in proteins that function in glutamate reuptake. Disrupting glutamate reuptake could elevate its concentration in the synapse and cause smaller changes in glutamate concentration in responses to stimuli. In this regard the more depolarized resting membrane potential of the BC1s (that we report) in the *Lrit3*^−^*^/^*^−^ retinas supports this conclusion.

In conclusion, our studies show that LRIT3 not only controls the post-synaptic assembly of the signaling complexes in DBCs, but also disrupts glutamate release or handling in the cone synaptic cleft resulting in decreases in excitatory responses in OFF BCs, and thus OFF RGCs.

## Extended Figure Legend

**Figure 2–1.**
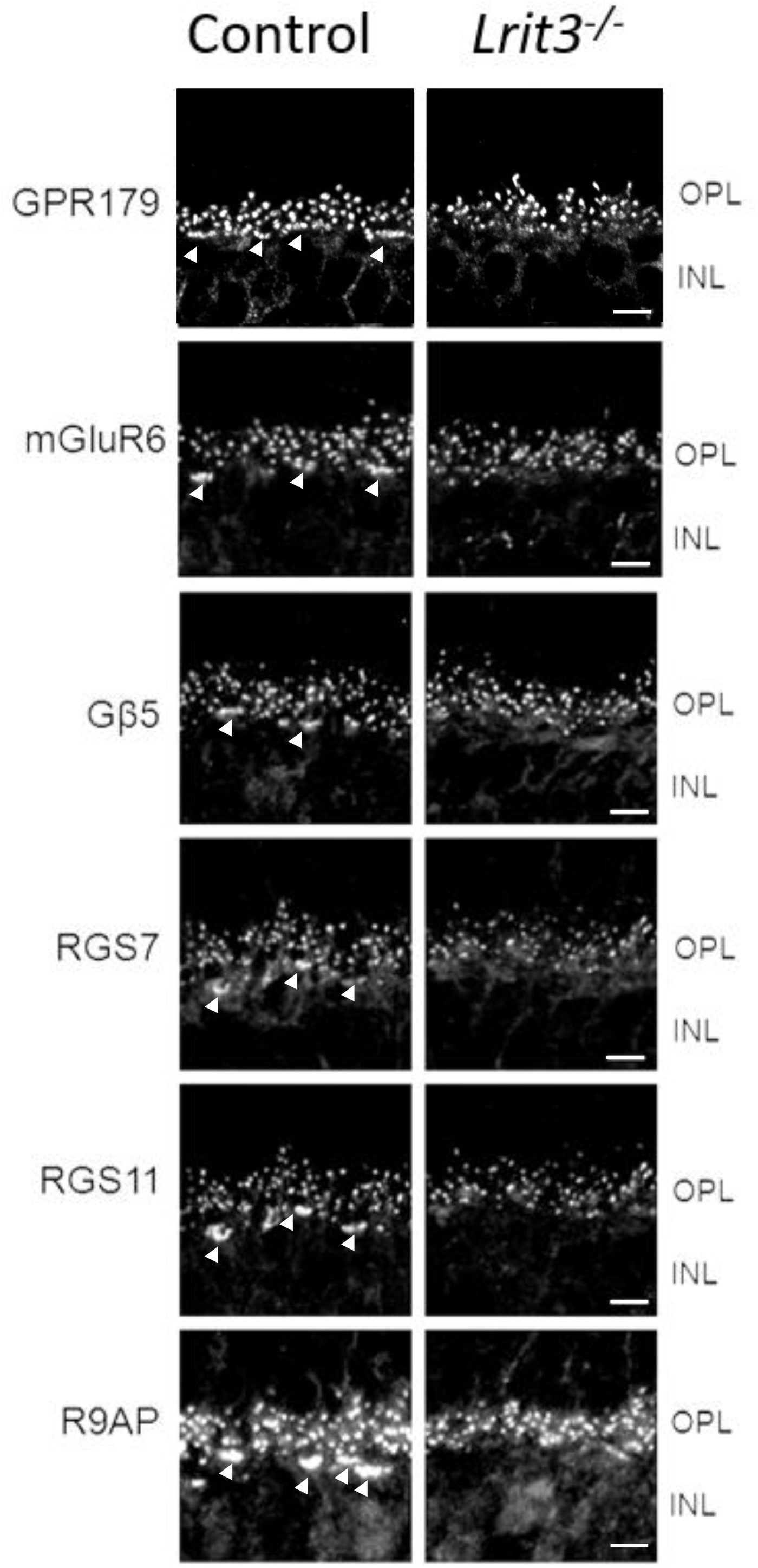
The absence of LRIT3 has differential effects on rod and cone DBC signalplexes. Immunohistochemical staining for GPR179, mGluR6, Gβ5, RGS7, RGS11 and R9AP show punctate staining at the dendritic tips of both rod and cone (large clusters at the base of the OPL and indicated by arrowheads) DBCs in control mice. In *Lrit3*^−^*^/^*^−^ mice these proteins are localized to the rod DBC dendritic tips but are absent from the cone DBCs. Note the lack of the large clusters at the base of the OPL. Scale bar = 5μm. OPL, outer plexiform layer; INL, inner nuclear layer.

## Acknowledgements

This work was supported by funding from the National Institutes of Health (R01 EY12354 (RGG, MAM); R01 EY014701 (MAM), R01 EY028188 (BGB) and an unrestricted grant from Research to Prevent Blindness to the University of Louisville.

